# Early life stress in male mice blunts responsiveness in a translationally-relevant reward task

**DOI:** 10.1101/2023.03.20.533443

**Authors:** Erin E. Hisey, Emma L. Fritsch, Kerry J. Ressler, Brian D. Kangas, William A. Carlezon

## Abstract

Early-life stress (ELS) leaves signatures upon the brain that persist throughout the lifespan and increase the risk of psychiatric illnesses including mood and anxiety disorders. In humans, myriad forms of ELS—including childhood abuse, bullying, poverty, and trauma—are increasingly prevalent. Understanding the signs of ELS, including those associated with psychiatric illness, will enable improved treatment and prevention. Here we developed a novel procedure to model human ELS in mice and identify translationally-relevant biomarkers of mood and anxiety disorders. We exposed male mice (C57BL/6J) to an early-life (juvenile) chronic social defeat stress (jCSDS) and examined social interaction and responsivity to reward during adulthood. As expected, jCSDS-exposed mice showed a socially avoidant phenotype in open-field social interaction tests. However, sucrose preference tests failed to demonstrate ELS-induced reductions in choice for the sweetened solution, suggesting no effect on reward function. To explore whether other tasks might be more sensitive to changes in motivation, we tested the mice in the Probabilistic Reward Task (PRT), a procedure often used in humans to study reward learning deficits associated with depressive illness. In a touchscreen PRT variant that was reverse-translated to maximize alignment with the version used in human subjects, mice exposed to jCSDS displayed significant reductions in the tendency to develop response biases for more richly-rewarded stimuli, a hallmark sign of depression (anhedonia) when seen in humans. Our findings suggest that translationally-relevant procedures that utilize the same endpoints across species may enable the development of improved model systems that more accurately predict outcomes in humans.

## INTRODUCTION

Early life stress (ELS) is a primary risk factor for developing mood and anxiety disorders including Major Depressive Disorder (MDD), Generalized Anxiety Disorder (GAD), and Post-Traumatic Stress Disorder (PTSD)^1,2^. In humans, ELS—which encompasses many conditions including child abuse, bullying, neglect, and poverty/low resource environments—affects as many as 1 in 7 children a year and is increasingly associated with alterations in brain connectivity and anatomy during adolescence and adulthood^3–5^. Interestingly, studies in laboratory animals indicate that different forms of ELS can produce profoundly different neurobiological alterations. As only one example, early life neglect can result in precocious maturation of prefrontal amygdala circuitry, whereas early life physical abuse instead delays maturation of this same circuitry^6,7^. Considering that the effects of ELS on the brain seem so dependent on the specific attributes of the stressful experience, it is critical to explore multiple forms of ELS in model systems that are sensitive to key diagnostic features of mood and anxiety disorders, such as social withdrawal and decreased sensitivity to reward (anhedonia).

While numerous rodent models of early life neglect have been developed^8,9^, there are relatively few reports describing the effects of early life trauma with a physical component^10–12^. Chronic social defeat stress (CSDS) in adult rodents is a commonly used model of stress that produces deficits in endpoints including social interaction, motivation, and sleep^13–16^. Complex behavioral tasks such as intracranial self-stimulation (ICSS) clearly demonstrate that CSDS produces anhedonia^17^, but that procedure requires surgical intervention, long periods of training (weeks to months), and a substantial equipment infrastructure while having no complementary paradigm in humans to validate direct translational relevance. Instead, the sucrose preference test (SPT) is widely used to quantify hedonic motivation^18,19^, as it is rapid (days), inexpensive, and produces outcomes that generally align with data using other testing procedures. Indeed, CSDS in mice causes reliable decreases in preference for sucrose solutions in the SPT, a putative reflection of stress-induced anhedonia and depressive-like behavior^18,19^.

Here we describe the development of a novel juvenile chronic social defeat stress (jCSDS) procedure that produces profound and persistent changes in behavior during adulthood as the result of a single epoch of stress (10 days of CSDS) that occurs early in life. Whereas most reports on CSDS involve both stress exposure and testing during adulthood^20^, this new approach involves stress exposure in young mice and testing during adulthood. Specifically, four week-old C57BL/6 male mice are defeated by novel aggressive adult males of a different strain (Swiss Webster [CFW]) for 10 consecutive days. After each defeat session, the mouse pair is housed together in the same cage but separated by a plastic barrier. We report that, as expected, mice defeated as juveniles subsequently show robust social avoidance in tests conducted during adulthood. Surprisingly, however, behavior in the SPT^18,21^ is unaffected. To explore whether other tasks might be more sensitive to changes in motivation, we tested jCSDS-exposed mice in the Probabilistic Reward Task (PRT), a procedure often used in humans to study reward learning deficits associated with depressive illness. In a touchscreen version of the PRT that was directly reverse-translated from that used in humans and requires a modest training period (days to weeks), we found that jCSDS reduces response bias for richly rewarded trials, without producing alterations in task acquisition rates, accuracy, or reaction time. We also explored ^18,21^relationships among behavior in the social interaction tests, SPT, and PRT. Our findings are consistent with RDoC frameworks^22^ that emphasize heterogeneity of neuropsychiatric disorders via subdomains that may be differentially affected and dependent on the type of ELS experienced, while also highlighting the interpretational advantages of using translationally-relevant procedures and endpoints.

## METHODS

### Subjects

Three week-old male C57BL/6J (C57) mice were obtained from Jackson Laboratory (Bar Harbor, ME, USA). The mice were housed under a 12-hour light and 12-hour dark schedule temperature (21 ±2 C and humidity (50 ±20%), and food and water were available *ad libitum* until PRT training. Following arrival at McLean Hospital, the mice were given seven days to habituate to the vivarium prior to starting the experiment. All experimental procedures were approved by the Institutional Animal Care and Use Committee at McLean Hospital and were performed in accordance with the National Institutes of Health’s (NIH) Guide for the Care and Use of Animals.

### Juvenile chronic social defeat stress (jCSDS)

Virgin male CFW mice (8 weeks, Charles River) were housed with ovariectomized female CFW (8 weeks, Charles River) mice for at least 1 week and screened for aggression with C57 male mice prior to defeat sessions. Only male CFWs that show an attack latency of less than 30 seconds for 2 consecutive days are used for defeats. Before the start of the first defeat, female CFW mice were permanently removed from the cage.

Male C57 mice (P29 ±3 days) were exposed to a traditional 10-day defeat paradigm^20^, with slight modifications. Specifically, C57 juveniles were placed into the home cage of an aggressive male CFW mouse. Each session consists of physical interactions that proceed until the aggressor has delivered 30 bites. Defeat sessions end after 5 minutes maximum if fewer than 30 bites occur. The juvenile mice are then housed side-by-side with the male CFW aggressor overnight using a plastic cage divider that physically separates the mice but enables visual and olfactory contact. This procedure was repeated each day with a novel aggressor for 10 consecutive days. At 24 hours after the last defeat, each defeated C57 mice was re-housed with another defeated mouse but separated by plastic divider, to control for potential effects of social isolation in adolescence. Control mice were housed with other control mice but separated by a plastic divider from P29 onward, to ensure consistent housing conditions across conditions.

### Open Field Social Interaction (OFSI)

Mice were transported from the animal care facility and habituated to a behavioral testing room for 1 hour. After habituation, each C57 mouse was placed in a large square arena containing an empty wire cup for 150 seconds. The C57 was then removed and a non-aggressive CFW male was placed under the wire cup. The C57 was then placed back in the arena for an additional 150 seconds. Social interaction ratio was calculated as the amount of time spent in the ‘social interaction zone’ (~2 cm circular zone around the wire cup) when the CFW was inside the cup divided by the amount of time spent in the ‘social interaction zone’ when the cup was empty.

### Sucrose preference test (SPT)

Preference for a weak sucrose solution (1% wt/vol) to water was measured using a 3-day sucrose preference test via a two-bottle choice paradigm, as described^18^. First, mice were acclimated for two days with access to two bottles containing water. On Day 3, mice were given free access to one bottle containing water and the second containing the sucrose solution. Bottles were weighed every 24 hours and alternated sides to avoid a side-bias. Sucrose preference scores were calculated as the ratios of sucrose intake to total volume intake.

### Probabilistic reward task (PRT)

#### PRT training

The mouse version of the PRT is a touchscreen task reverse-engineered from human studies^23^ and modified from a variant originally designed for rats to objectively quantify reward responsivity^24,25^. Three days before PRT training commenced, mice were food-restricted to 80-85% of their original body weights via post-session portions of 2-3 grams of rodent chow, followed by training as described previously^25^. Briefly, mice were trained to rear and touch a 5×5 cm blue square on a black background and in various positions on the touchscreen to receive 0.02 mL of a highly palatable 20% sweetened condensed milk reward, which was delivered in a well on the opposite wall of the touchscreen. After reliable responding was observed, mice were then trained during 100-trial sessions to discriminate between a long or short white line (24×3cm or 12×3cm) on a black background by responding on one of two virtual levers (5×5cm blue squares) presented below the line, to the left and right of center. Correct responses were rewarded with sweetened condensed milk paired with a tone and brief, bright yellow screen followed by a 10 s blackout period, whereas incorrect responses resulted in a 20 s timeout. During initial PRT training sessions, a correction procedure^26^ was employed, where incorrect trials were repeated until a correct response was made prior to advancing to the next trial. After reaching criterion in this phase (10 or fewer errors for both the long and short lines on two consecutive sessions), mice were tested without correction under otherwise identical contingencies. Once mice reached >80% accuracy during two consecutive sessions without correction, PRT testing commenced.

#### PRT testing

Following line-length discrimination training to criteria, subjects were exposed to a 5-session PRT testing protocol using 3:1 probabilistic reinforcement contingencies such that a correct response to one of the line lengths (long or short) was reinforced 60% of the time (rich stimulus), whereas a correct response to the other line length was reinforced 20% of the time (lean stimulus). Incorrect responses were never reinforced. The line length associated with the rich and lean contingency was determined for each subject during their final two line length discrimination training sessions by examining their accuracies and designating the line length with a higher mean accuracy as the stimulus to be rewarded on the lean schedule. This approach was expressly designed to examine the effects of jCSDS on response bias generated by responsivity to asymmetrical probabilistic contingencies, rather than the amplification of a preexisting inherent bias that is a function of uncontrolled variables.

### Statistical Analyses

#### PRT

The PRT yields two primary dependent measures: response bias (which reflects reward function) and discriminability (which reflects baseline response capabilities). These values can be quantified using equations derived from signal detection theory^24^ by examining the number of _Correct_ and _Incorrect_ responses for Rich and Lean trial types. Response Bias is calculated using the following *log b* equation:

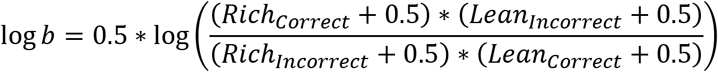

High bias values are produced by high numbers of correct responses for rich trials and incorrect responses for lean trials. Discriminability is calculated using the following *log d* equation:

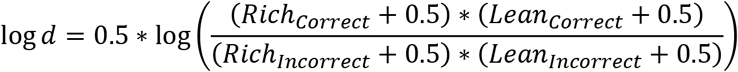

High discriminability values are produced by high numbers of correct responses for both rich and lean trials. (0.5 is added to all parameters in both equations to avoid instances where no errors are made on a given trial type, thus making log transforms impossible.) In addition, reaction time (latency from line presentation to response) was calculated and presented as individual subject values and session-wide group means (±SEM) for rich and lean trials.

#### OFSI, SPT, and PRT

Unpaired t-tests were used to evaluate group differences (control vs. jCSDS) on behavior in each of the test procedures. A two-way repeated measures ANOVA was used to evaluate differences in main effect between groups for log b and log d collected each day of PRT testing. The criterion for significance was set at *p*<0.05. All statistical analyses were conducted using GraphPad Prism 9 Software (San Diego, CA, USA).

## RESULTS

### Experiment 1

Experiment 1 was designed to characterize the effects of the novel jCSDS procedure on social interaction (using the OFSI test) and reward function (using the SPT) (**Fig. 1A**). Based on published work describing the effects of adult CSDS^18,19^, we hypothesized that we would observe stress-induced reductions in both of these metrics following jCSDS.

**Figure 1.**
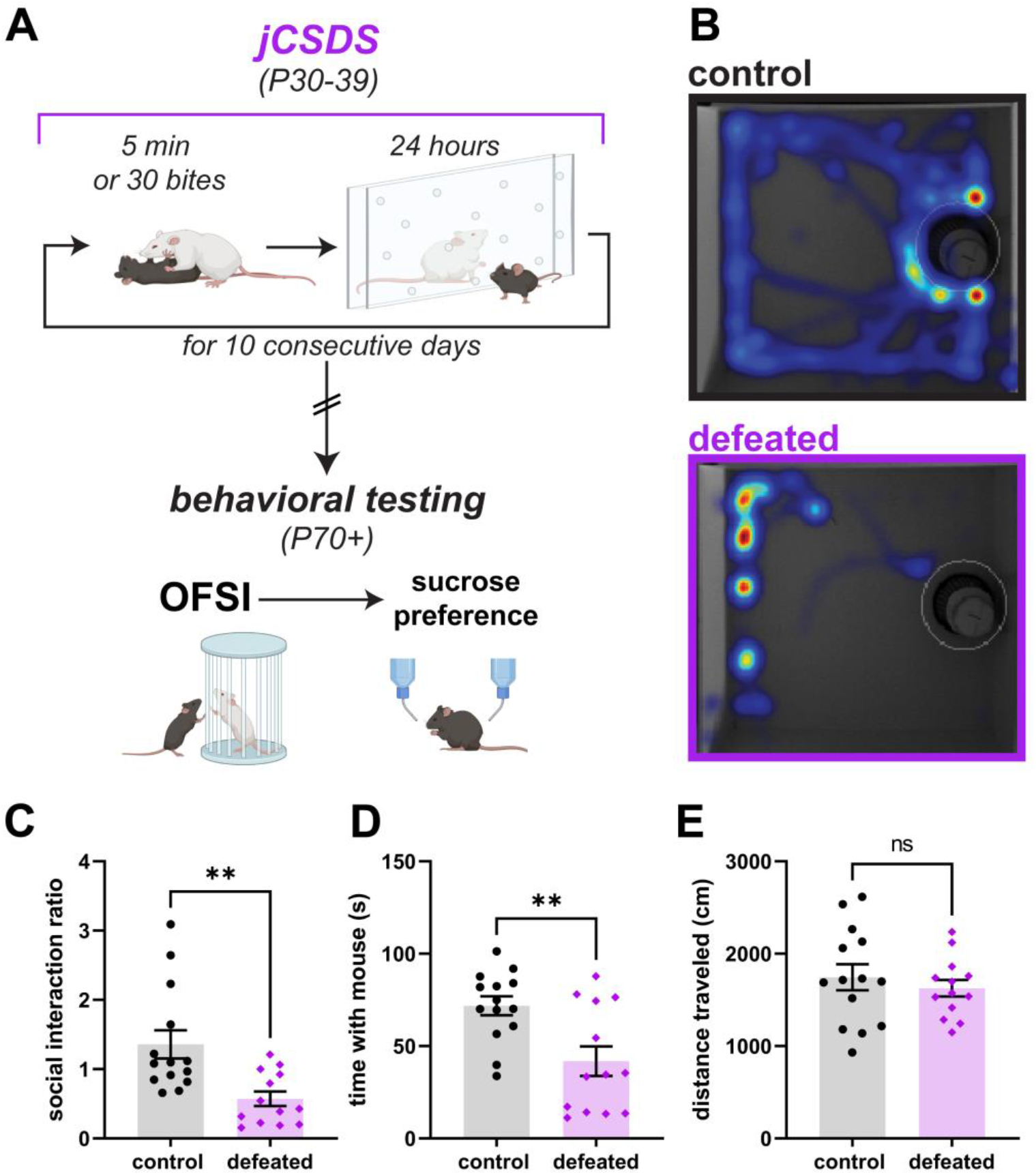
Social interaction is reduced in adults defeated as juveniles. (A) Schematic of jCSDS and behavioral experiments. B) Example heatmaps from open field social interaction test (OFSI). C) Social interaction ratio in control (black) versus defeated (purple) adults. D) Time spent interacting with non-aggressive CFW. E) Total distance traveled in centimeters over OFSI testing.

### Adults defeated as juveniles show reduced social interaction

Juvenile male C57BL/6J mice were subjected to the 10-day jCSDS regimen. Each defeated mouse was then housed in the same cage as another defeated mouse, separated with a plastic barrier that allows for olfactory and visual contact, for the duration of the experiment. Control mice were housed beside other control mice in the same manner to control for housing conditions. Once the mice reached adulthood (P60-70), we examined social approach to a non-familiar non-aggressive male CFW using in the OFSI (**Fig. 1B**). While control mice showed increased preference for social interaction (i.e., social interaction ratio >1 indicates more time in interaction zone when another mouse is present than when absent), mice exposed to jCSDS avoided social interaction, as evidenced by a low social interaction ratio (unpaired two-tailed t-test, n=14 controls, 13 defeated, t[df]=3.350[25], p=0.003) (**Fig. 1C**) as well as a significantly reduced raw social interaction time compared to controls (t[df]=3.209[25], p=0.004) (**Fig. 1D)**. There were no differences in distance traveled (t[df]=0.696[25], p=0.493) (**Fig. 1E**), suggesting that the change in social interaction was not simply a result of changes in locomotor activity. These findings suggest that defeated mice retain a long-lasting memory of juvenile trauma into adulthood, creating strong signs of social avoidance behavior.

### Adults defeated as juveniles show no difference in sucrose preference

As depressive signs during adulthood is often comorbid with early life trauma in humans, we next wanted to examine depressive-like behaviors in adult mice defeated as juveniles to determine if our model of early life trauma could recapitulate the signs of anhedonia. Following the OFSI tests, we tested the mice in the SPT, which has been used extensively in studies of adult CSDS^27^. Surprisingly, defeated mice showed no difference in sucrose preference for a 1% sucrose solution (**Fig. 2A**) (unpaired two-tailed t-test, n=14 control, 13 defeated, t[df]=0.082[25], p=0.940). Interestingly, defeated mice showed an increase in total volume of liquid consumed (t[df]=2.883[25], p=0.008) (**Fig. 2B**) and amount of sucrose consumed per bodyweight (t[df]=2.415[25] p=0.023) (**Fig. 2C**). These findings suggest that juvenile exposure to stress does not produce anhedonia—a finding inconsistent with a vast human literature documenting the persistent effects of ELS across the lifespan—or alternatively, that the SPT is not sensitive to the specific types of reward deficits that are produced by jCSDS.

**Figure 2.**
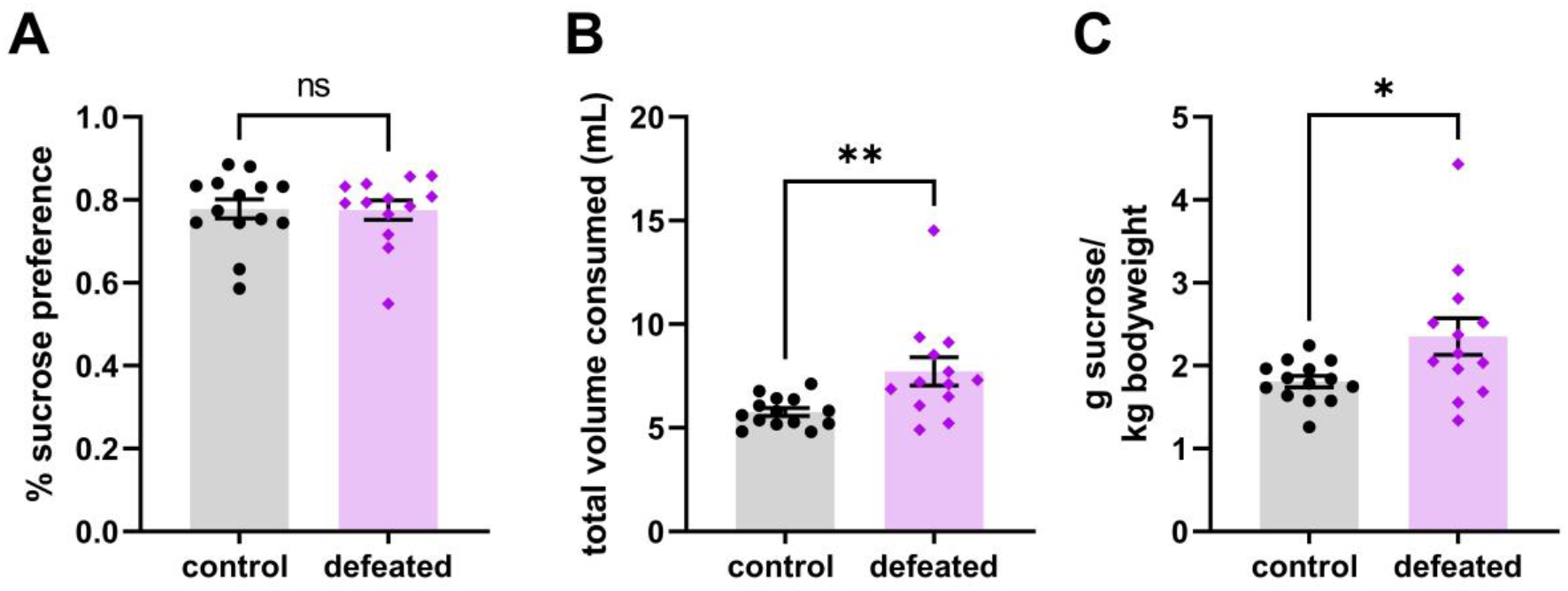
Sucrose preference is unaffected in adults defeated as juveniles. Average percent sucrose preference across 3 day testing period (A). Average total volume of sucrose solution and water consumed per day calculated across 3 day testing period (B). Average sucrose consumed per body weight per day calculated across 3 day testing period (C).

### Experiment 2

Experiment 2 was designed to determine if jCSDS effects would be detectable in the PRT, a task frequently used to study reward function in humans with mood and anxiety disorders^27^. For consistency with Experiment 1, new cohorts of mice were first tested in the OFSI, followed by the PRT instead of the SPT (**Fig. 3A**). After the PRT, the mice were tested in the SPT, to explore the reproducibility of the findings (lack of effect) from Experiment 1. Based on published work describing the performance of humans with depressive illness in the PRT^24^, we hypothesized that this task may identify anhedonia-like signs in mice exposed to jCSDS.

**Figure 3.**
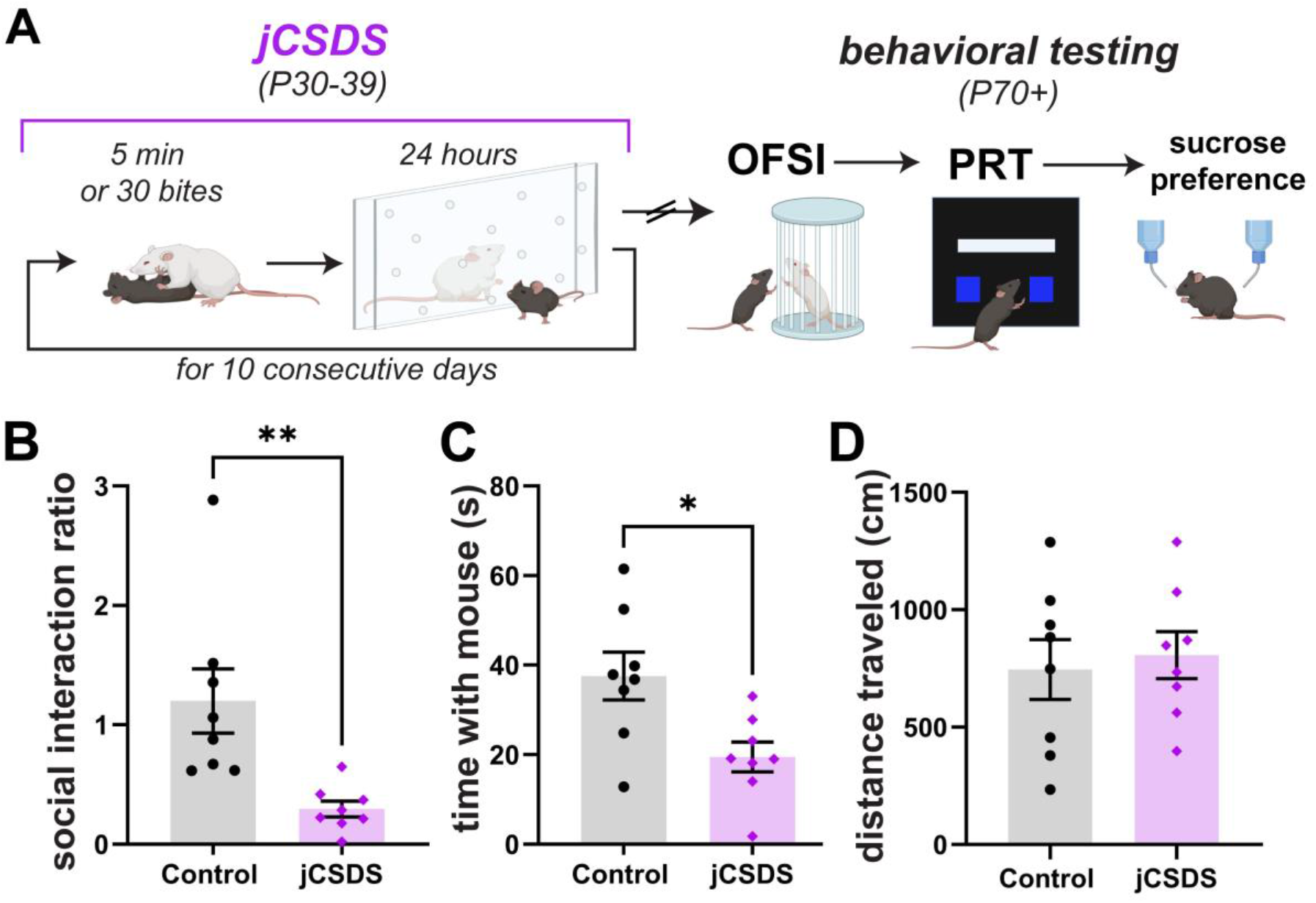
Replication of social interaction deficits in new cohort of adults defeated as juveniles. (A) Schematic of jCSDS and behavioral experiments. B) Social interaction ratio in control (black) versus defeated (purple) adults. C) Time spent interacting with non-aggressive CFW. D) Total distance traveled in centimeters over OFSI testing.

### Replication: adults defeated as juveniles show reduced social interaction

As was the case in Experiment 1, while control mice showed increased preference for social interaction, mice exposed to jCSDS avoided social interaction, as indicated by a low social interaction ratio (n=8 controls, 8 defeated, t[df]=3.273[14], p=0.006, 1 defeated mouse was excluded from social interaction analysis as an outlier (Grubb’s outlier test, alpha=0.05)) (**Fig. 3B)** as well as a significantly reduced raw social interaction time compared to controls (t[df]=2.889[14], p=0.012) (**Fig. 3C)**. There were no changes in distance traveled (t[df]=0.375[14], p=0.713) (**Fig. 3D**), suggesting that the change in social interaction was not simply a result of changes in locomotor activity. These findings replicate the OFSI findings from Experiment 1 and suggest that the effect is rigorous and reproducible across independent cohorts of mice exposed to jCSDS.

### Adults defeated as juveniles show blunted response bias in the PRT

PRT procedures were initiated at P90-120. Defeated mice showed no difference in learning rates in any of the training stages of PRT compared to control mice (**Supplemental Fig. 1A-D**). After reaching line-length discrimination training criterion (**Fig. 4A**), mice were tested on the PRT for 5 daily sessions. Compared to controls, mice exposed to jCSDS showed profoundly blunted response biases towards more richly rewarded stimulus (log b; n=8 controls, 9 defeated; unpaired two-tailed t-test, t[df]=2.978[15], p=0.009) (**Fig 4B**), although both groups performed with similar levels of task discriminability (log d; t[df]=1.382[15], p=0.187) (**Fig. 4C**). This blunted response bias could not be accounted for by differences in reaction time (t[df]=0.293[15], p=0.773) (**Fig. 4D**) or body weight (t[df]=0.051[15], p=0.960) (**Fig. 4E**). Whereas both defeated and control mice showed steady increases in response bias over the course of the 5-day PRT regimen, it was consistently lower in the jCSDS-exposed mice (2-way repeated measures ANOVA, main effect of defeat status: F[1,15]=9.050, p=0.009) (**Supplemental Fig. 2A**). In contrast, whereas both defeated and control mice showed comparable increases in discriminability over the same period, there were no group differences at any time point (main effect of defeat status: F[1,15]=1.939, p=0.184) (**Supplemental Fig. 2B**). These findings suggest that jCSDS in mice causes depressive-like effects that persist into adulthood, and that the PRT is more sensitive to this outcome than the SPT.

**Figure 4.**
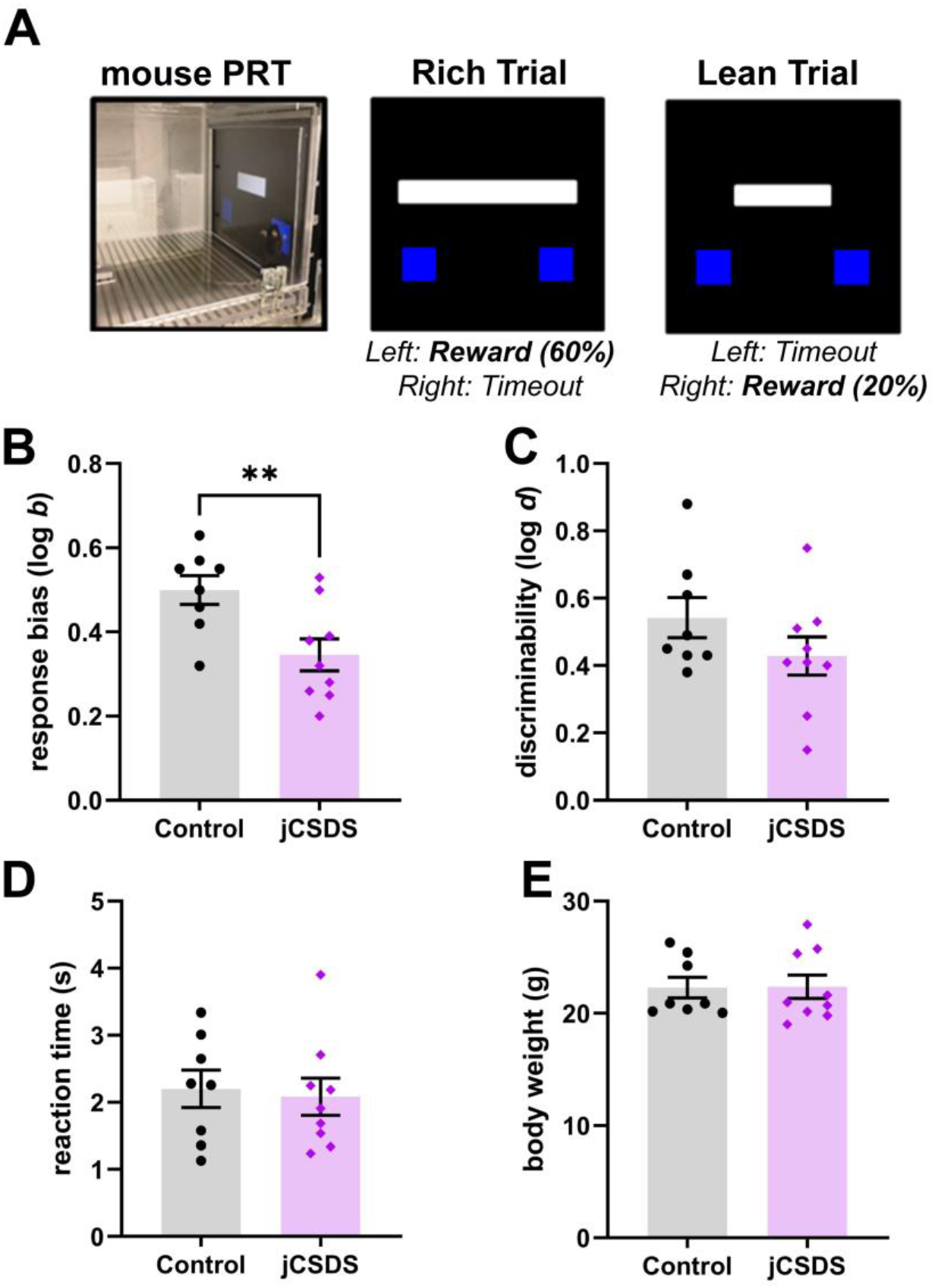
Response bias is reduced in adults defeated as juveniles. A) Schematic of Probabilistic Reward Task (PRT). B) Averaged response bias (log b) over 5 days of PRT testing. C) Averaged discriminability (log b) in control (black, n=8) and adults defeated as juveniles (jCSDS, purple, n=9) over 5 days of PRT testing. (D) Averaged reaction time in seconds over 5 days of PRT testing. E) Body weight on the last day of PRT testing.

### Replication: adults defeated as juveniles show no differences in sucrose preference

After PRT testing, mice were taken off food restriction and given food and water ad libitum for 1 to 2 weeks before the SPT (P150-180). Despite the different order of testing and an intervening period of food restriction, as was the case in Experiment 1, defeated mice showed no difference in sucrose preference for a 1% sucrose solution (**Fig. 5A**) (n=8 control, 9 defeated, t[df]=0.636[15], p=0.534). Defeated mice also showed no changes in total volume of liquid consumed (t[df]=0.735[15], p=0.474) (**Fig. 5B**) or amount of sucrose consumed per bodyweight (t[df]=0.358[15], p=0.726) (**Fig. 5C**). These findings replicate the SPT findings from Experiment 1, albeit under slightly different order of testing procedures, and suggest that the lack of effect in this behavioral paradigm is rigorous and reproducible across independent cohorts of mice exposed to jCSDS.

**Figure 5.**
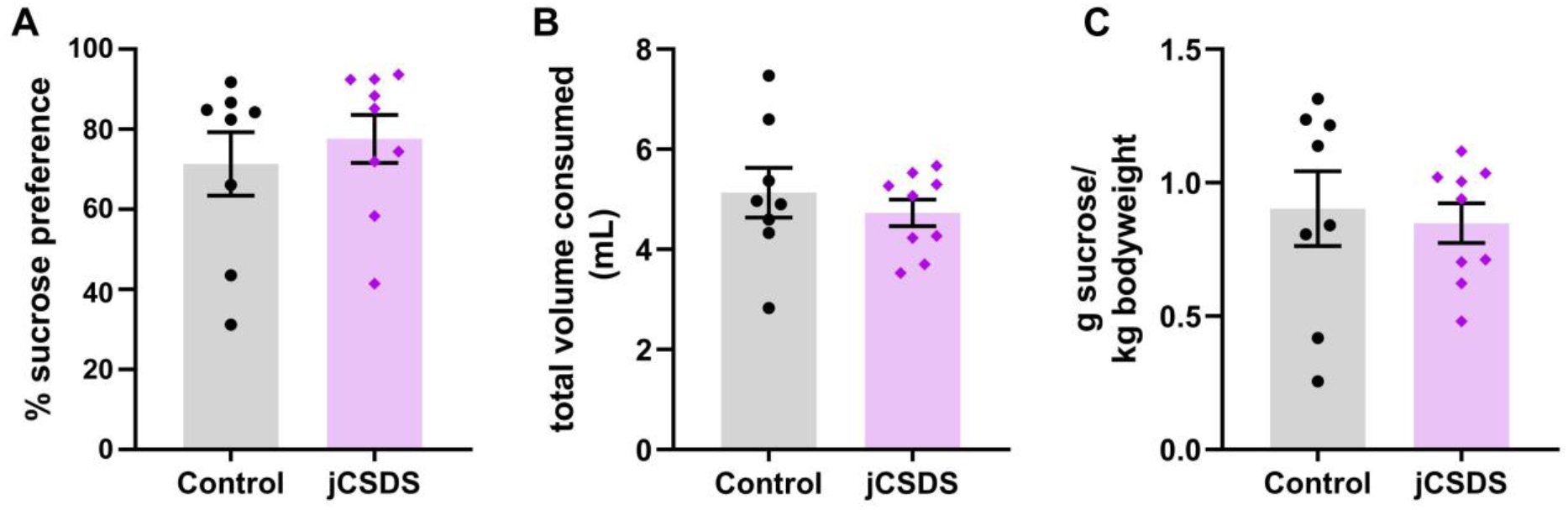
Replication of unaffected sucrose preference in adults defeated as juveniles. Average percent sucrose preference across 3 day testing period (A). Average total volume of sucrose solution and water consumed per day calculated across 3 day testing period (B). Average sucrose consumed per body weight per day calculated across 3 day testing period (C).

### Associations among behaviors in the OFSI, SPT, and PRT

Using the data from mice tested in Experiment 2, we next examined if the behavioral endpoints in any of the tests correlated with one another. Interestingly, percentage sucrose preference was unrelated to either social interaction ratio (Pearson correlation coefficient, R=0.108, p=0.680) (**Supplemental Fig. 3A**) or response bias (Pearson correlation coefficient, R=-0.067, p=0.805) (**Supplemental Fig. 3B**). Social interaction ratio and response bias showed a nominal, though not statistically significant, correlation (Pearson correlation coefficient, R=0.388, p=0.138) (**Supplemental Fig. 3C**). These data are consistent with the conclusion that these endpoints reflect separate behavioral domains that are not necessarily coupled to one another.

## DISCUSSION

Here we demonstrate that jCSDS a novel variant of prototypical CSDS procedures in which juvenile (instead of adult) mice are exposed to stress and tested during adulthood, produces robust and persistent changes in behavior. We focused on social interaction and reward function, which are two RDoC-related domains (social processes, positive valence)^22^ that are often dysregulated in mood and anxiety disorders^28^. First, jCSDS causes robust decreases in social behavior (i.e., social avoidance) that are unrelated to changes in locomotor activity. Additionally, jCSDS produces a blunted response bias to richly rewarded trials in PRT, an outcome reflecting anhedonia in humans and rodents^29^. Surprisingly, however, jCSDS failed to alter sucrose preference, an assay commonly used to quantify hedonic behavior in rodents. The observation that jCSDS causes social avoidance but no change in sucrose preference was replicated in two independent cohorts of mice, indicating that it is rigorous and reproducible. Importantly, the SPT was conducted at two different times in the cohorts—immediately after OFSI tests in the first cohort and after OFSI and PRT testing in the second cohort—making it seem unlikely that the failure to observe anhedonia is related to not allowing sufficient time for the phenotype to develop. This pattern of results suggests that the PRT is more sensitive than the SPT to stress-induced changes in reward function, or alternatively, that it may reveal additional facets of anhedonia that are not captured by the SPT alone. Regardless, this work in mice is consistent with a large literature indicating that early life physical and emotional trauma in humans can cause profound changes in the behavior and brain biology that persist across the lifespan^4,5^ and thus establishes a novel methodology that may be useful for modeling ELS-related psychiatric illness.

It is important to note that anhedonia is a construct characterized by substantial heterogeneity and subdomains, and it is unlikely that either the SPT or the PRT captures its full spectrum. Indeed, these tests are expressly designed to evaluate different aspects of reward responsiveness (reward consumption versus reward learning, respectively). The lack of concordance or correlation between assay outcomes, which has been observed previously^28^, is both not surprising and consistent with an RDoC-like framework that requires a diversity of tasks to investigate these multidimensional psychological systems. While it is conceivable that subtle changes in behavior in the SPT may have been detectable if a full range of sucrose concentrations had been examined—analogous to dose-finding in drug self-administration studies—a null effect is more easily interpreted than reductions in intake (which could reflect either leftwards or rightward shifts in functions represented by an inverted U-shaped curve). The lack of correlations among the behavioral endpoints in the OFSI, SPT, and PRT suggest that social processes and positive valence can be uncoupled in mice, as is the case in humans, where certain signs are frequently co-morbid but there is considerable heterogeneity among patients and myriad combinations among conditions^28^.

The approaches used here may help to promote improvements in the face validity and translational relevance of preclinical studies of human conditions the enhance risk of developing mental health disorders. While ELS can take many forms in humans, jCSDS may most closely approximate bullying, considering that the subordinate mice are younger, smaller, and not genetically related to the aggressors. The mouse version of the PRT was reverse-translated from a version used in humans and involves a digital element (responses delivered via touchscreen), which together enhance the translational relevance of the work. In humans, significant reductions in the tendency to develop response biases for more richly-rewarded stimuli is frequently seen with depressive illness and considered a hallmark sign of anhedonia^23^. The fact that the PRT has been so thoroughly validated in humans and the mouse version shares conceptual, procedural, and analytical elements highlights the interpretational advantages of using translationally-relevant procedures and endpoints. Widespread adoption of these principles may enable the development of improved animal models that more accurately predict outcomes in humans^30^.

One limitation of the current experiments is that only male mice were studied. It is well-established that biological sex affects resiliency and susceptibility to stress. Indeed, the diagnoses of mood disorders in which anhedonia is prominent are more prevalent in women^31^. While our recent work with adult CSDS demonstrates that, with at least some stress biomarkers, effects in male mice accurately predicts outcomes in both traumatized men and women^32^, it is known that ELS in rodents can produce sex-dependent effects on reward^33^. Application of jCSDS to female mice is hypothetically feasible and currently under development in our labs, with efforts to develop a single standardized protocol that can be used for both sexes. Upon validation, future experiments will investigate sex differences in response to early life physical trauma.

In summary, we have demonstrated that the jCSDS model of ELS can recapitulate key symptoms of childhood trauma and comorbid psychiatric conditions. In male mice tested during adulthood, jCSDS produces a distinct socially avoidant and anhedonic phenotype. These findings provide the behavioral basis for investigations into the neural circuit alterations as well as more holistic (e.g., peripheral) biomarkers of early life trauma and abuse that may enable improvements in the diagnosis, treatment, and prevention of psychiatric illness.

---

**Supplemental Figure 1.**
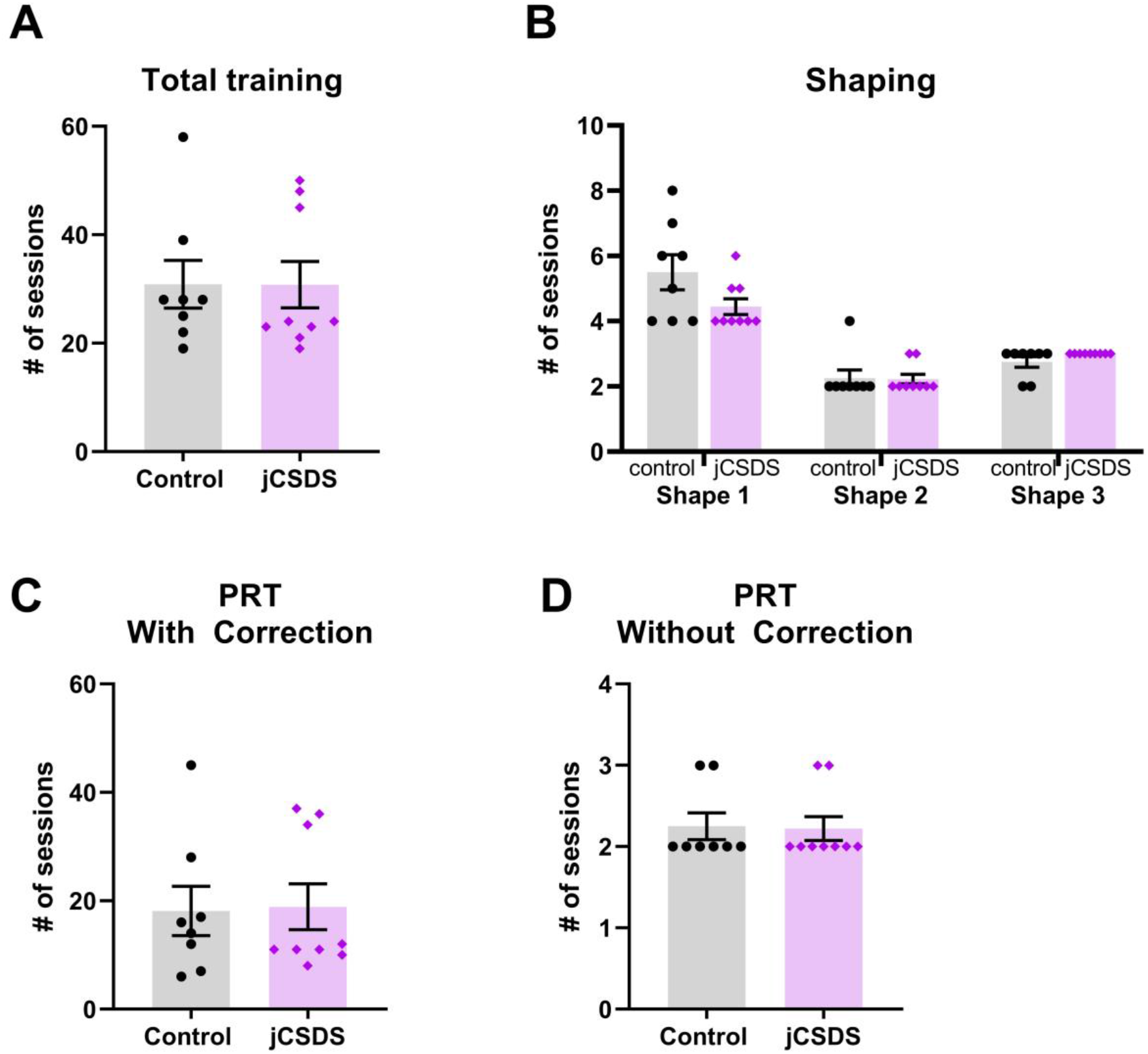
Number of sessions to completion for total and individual training phases is similar for control and defeated animals. A) Total number of training sessions to reach criteria for PRT testing for control (black) and defeated (purple) animals. B) Number of sessions to reach criteria for initial shaping (B), corrected (C), and uncorrected (D) phases of PRT training.

**Supplemental Figure 2.**
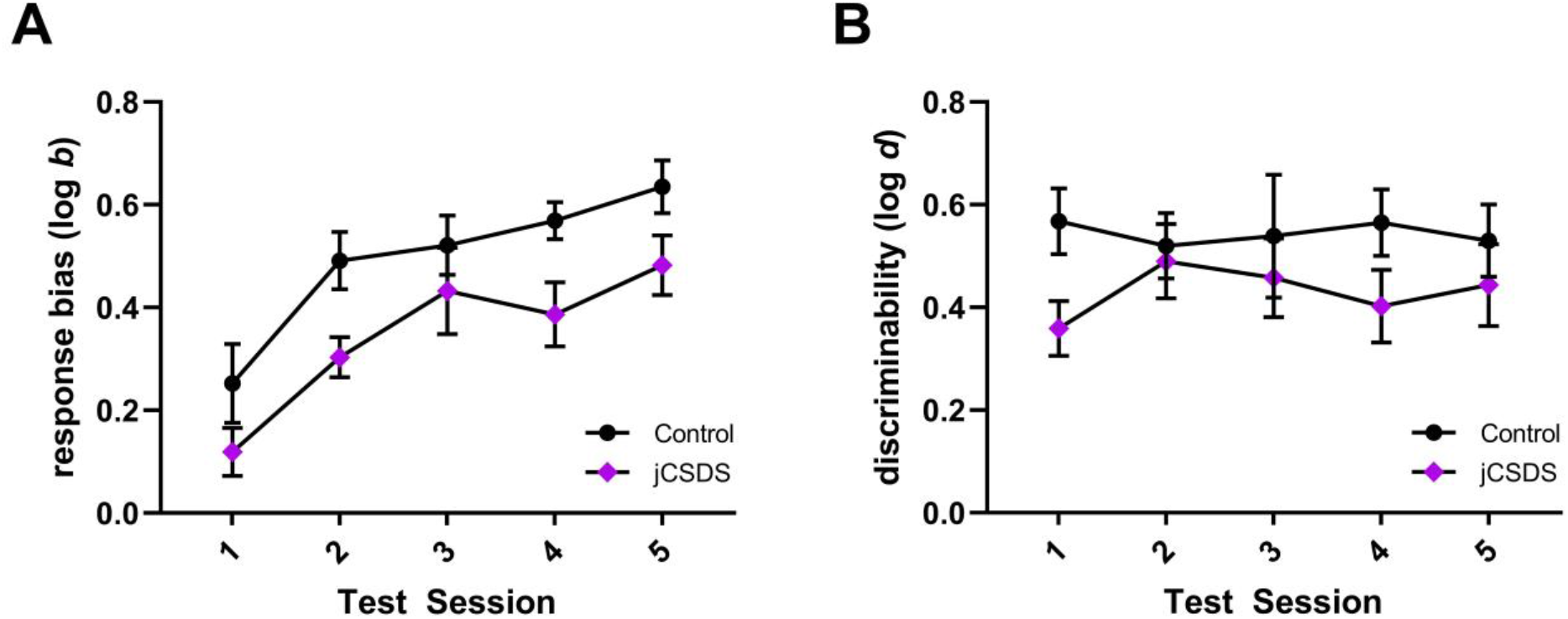
Discriminability and response bias values on each day of PRT testing. Response bias (log *b*, A) and discriminability (log *d*, B) and on each of the five days of PRT testing.

**Supplemental Figure 3.**
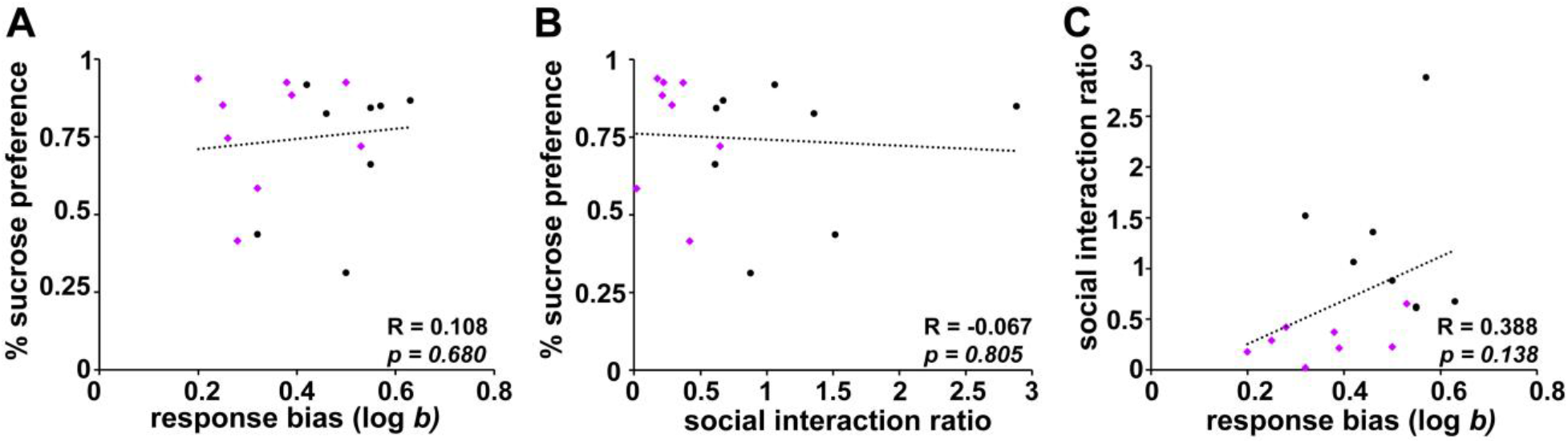
Correlations between response bias, sucrose preference, and social interaction. Correlations between percent sucrose preference as a function of response bias (A) or social interaction ratio (B) as well as social interaction as a function of response bias (C). Dotted lines indicate linear fit to control and defeated data. Pearson correlation coefficient (R) listed for combined control and defeated data.

